# Machine learning prediction and phyloanatomic modeling of viral neuroadaptive signatures in the macaque model of HIV-mediated neuropathology

**DOI:** 10.1101/2022.06.17.496109

**Authors:** Andrea S. Ramirez-Mata, David Ostrov, Marco Salemi, Simone Marini, Brittany Rife Magalis

## Abstract

In human immunodeficiency virus (HIV) infection, virus replication in the central nervous system (CNS) can result in HIV-associated neurocognitive deficits in approximately 25% of patients with unsuppressed viremia and is thought to be characterized by evolutionary adaptation to this unique microenvironment. While no single mutation can be agreed upon as distinguishing the neuroadapted population from virus in patients without neuropathology, earlier studies have demonstrated that a machine learning (ML) approach could be applied to identify a collection of mutational signatures within the envelope glycoprotein (Env Gp120) predictive of disease. The S[imian] IV-infected macaque is a widely used animal model of HIV neuropathology, allowing in-depth tissue sampling infeasible for human patients. Yet, translational impact of the ML approach within the context of the macaque model has not been tested, much less the capacity for early prediction in other, non-invasive tissues. We applied the previously described ML approach to prediction of SIV-mediated encephalitis (SIVE) using *gp120* sequences obtained from the CNS of animals with and without SIVE with 73% accuracy. The presence of SIVE signatures at earlier time points of infection in non-CNS tissues in both SIVE and SIVnoE animals indicated these signatures cannot be used in a clinical setting. However, combined with protein structural mapping and statistical phylogenetic inference, results revealed common denominators associated with these signatures, including 2-acetamido-2-deoxy-beta-D-glucopyranose structural interactions and the infection of alveolar macrophages. Alveolar macrophages were demonstrated to harbor a relatively large proportion (35 – 100%) of SIVE-classified sequences and to be the phyloanatomic source of cranial virus in SIVE, but not SIVnoE animals. While this combined approach cannot distinguish the role of this cell population as an indicator of cellular tropism from a source of neuroadapted virus, it provides a key to understanding the function and evolution of the signatures identified as predictive of both HIV and SIV neuropathology.

**Author summary:** HIV-associated neurocognitive disorders remain prevalent among HIV-infected individuals, even in the era of potent antiretroviral therapy, and our understanding of the mechanisms involved in disease pathogenesis, such as virus evolution and adaptation, remains elusive. In this study, we expand on a machine learning method previously used to predict neurocognitive impairment in HIV-infected individuals to the macaque model of AIDS-related neuropathology in order to characterize its translatability and predictive capacity in other sampling tissues and time points. We identified four amino acid and/or biochemical signatures associated with disease that, similar to HIV, demonstrated a proclivity for proximity to aminoglycans in the protein structure. These signatures were not, however, isolated to specific points in time or even to the central nervous system, as they could be observed at low levels during initial infection and from various tissues, most prominently in the lungs. The spatiotemporal patterns observed limit the use of these signatures as an accurate prediction for neuropathogenesis prior to the onset of symptoms, though results from this study warrant further investigation into the role of these signatures, as well as lung tissue, in viral entry to and replication in the brain.

## Introduction

Human Immunodeficiency Virus (HIV) is capable of infecting the central nervous system (CNS) and significantly damaging neighboring neuronal cells, ultimately leading to neurocognitive impairment. Clinical manifestations of HIV-associated neurocognitive disease (HAND), collectively referred to as neuroAIDS, can include decreased attention and concentration, memory loss, reduced psychomotor ability and executive function, and tremors; with time, these manifestations can progress to incapacitating weakness, paraperesis, and severe disease in the form of dementia (1). Despite the success of antiretroviral therapies (ART) in countries where they are widely available, neuroAIDS continues to persist among persons living with HIV (PLWH). Though more commonly observed among PLWH that have progressed to end-stage acquired immunodeficiency syndrome (AIDS) (2), not all PLWH (approximately 25%) are diagnosed (3), suggesting a distinct viral- or immune-mediated mechanism associated with disease pathogenesis that has yet to be identified (4;5).

In order for infection of the brain to occur, the virus must enter by way of the highly selective blood-brain barrier and replicate in a subset of cells – perivascular macrophages and microglia – that are unique to this tissue (6). The CNS thus constitutes a distinctive microenvironment imparting measurable selective pressure(s) that can give rise to tissue-adapted viral variants (7;8; 9;10). Multiple studies have explored further aspects of this hypothesis and concluded that there exist HIV-1 envelope *(Env)* amino acid patterns, or signatures, unique to viral sequences found in the CNS (7; 11; 12; 13; 14). Such individual signatures, however, vary depending on the study, rendering results inconclusive as to a single, critical mutation and/or mutational region, associated with neuroadaptation. The controversy surrounding the existence of a neuroadaptive signature may be explained by failing to take into account the combined effect of multiple, non-contiguous amino acid residues and their biochemical properties (15).

Machine learning (ML) approaches have been more recently applied in genomic analysis to model the correlation between disease variants and clinical outcomes, capitalizing on the presence of multiple, combinatorial viral factors (16; 17; 18). Holman and Gabuzda (2012)(15) developed a ML pipeline paving the way for the use of HIV viral variants in predicting neuropathological outcome. Their study identified five statistically significant amino acid signatures that were capable of predicting with 75% accuracy the outcome of HIV-associated dementia (HAD). However, this study was limited to analysis of a small portion of the gene – the V3 loop and surrounding C2 and C3 regions – and the position of the signatures within the linear sequence (i.e., no protein structure context). Ogishi and Yotsuyanagi (2018) (19) also generated a ML prediction model for HAND from a comprehensive data set, similarly to Holman and Gabuzda taking into account only the C2V3C3 region, in which they used an iterative ML and step-wise feature reduction. With this methodology, they obtained accuracy of 100% in a hold-out testing sub-dataset and 95% using the entire dataset, reporting that only three genetic features were sufficient to predict HAND status regardless of sampled tissue. Owing to more general limitations of the use of human subjects, no conclusions could be drawn regarding how early this prediction could be made in time; nor whether these signatures could be found in other tissues that comprise the potential origins for neurovirulence and thus targets for prevention of CNS invasion (20).

The simian immunodeficiency virus (SIV)-infected macaque model can be used to recapitulate neuropathogenesis associated with HAND and provides a unique opportunity to acquire additional tissue samples at numerous time points throughout the course of infection and/ or treatment (21;22). Approximately 30% of SIV-infected *Rhesus* macaques that naturally progress to simian AIDS (i.e., without immune modulation) develop the pathological hallmark of neuroAIDS – SIV encephalitis (SIVE) – in 2-3 years. For a more rapid progression to neuroAIDS, a model has been developed in which a monoclonal antibody is used to deplete animals of CD8^+^lymphocytes (23). Both CD8^+^-depleted and CD8^+^non-depleted, or naturally progressing, models have been characterized in depth and represent well the neurological complication of neuroAIDS (7).

In this study, we explore the existence of SIV genetic and/or biochemical signatures predictive of HIV/SIV-mediated neuropathology using a ML approach similar to that of Holman and Gabuzda (15). In addition to characterizing the translational applicability of the results in HIV-infected individuals using protein structural data, we significantly expanded the earlier study by 1) using the full envelope *(env gp120)* region for model training and 2) applying the learned signatures to additional SIV sequences from a variety of tissues and time points, therefore exploring SIVE intra-host heterogeneity in terms of time and tissue trajectories.

## Results

The main objective of this study was to identify SIV protein signatures associated with, and predictive of AIDS-related neuropathology. Viral *env gp120* sequences used in model training were extracted from the CNS of animals originating from different cohorts (Table 1) and diagnosed at necropsy, upon experiencing AIDS-related symptoms, as SIVE/SIVnoE. First, we estimated the performance of the model by a leave-one-animal-out cross-validation. That is, we iteratively removed all the sequences from a single animal and used the as test set fold, while considering the rest as training. For each test fold of the cross-validation we classified both each single sequence and the animal as a whole (final animal classification as SIVE or SIVnoE if 85% sequences or more classified as either since this was the lowest percentage obtained for a given classification). The model outputs certain requirements, or “rules”, referred to as amino acid *signatures,* that sequences need to follow to be classified as SIVE or SIVnoE. Our final model was then appled to the sequences from non-CNS tissues at different collection time points during infection (application dataset) (Table 1) in order to evaluate spatiotemporal patterns of signature presence.

**Table 1.**
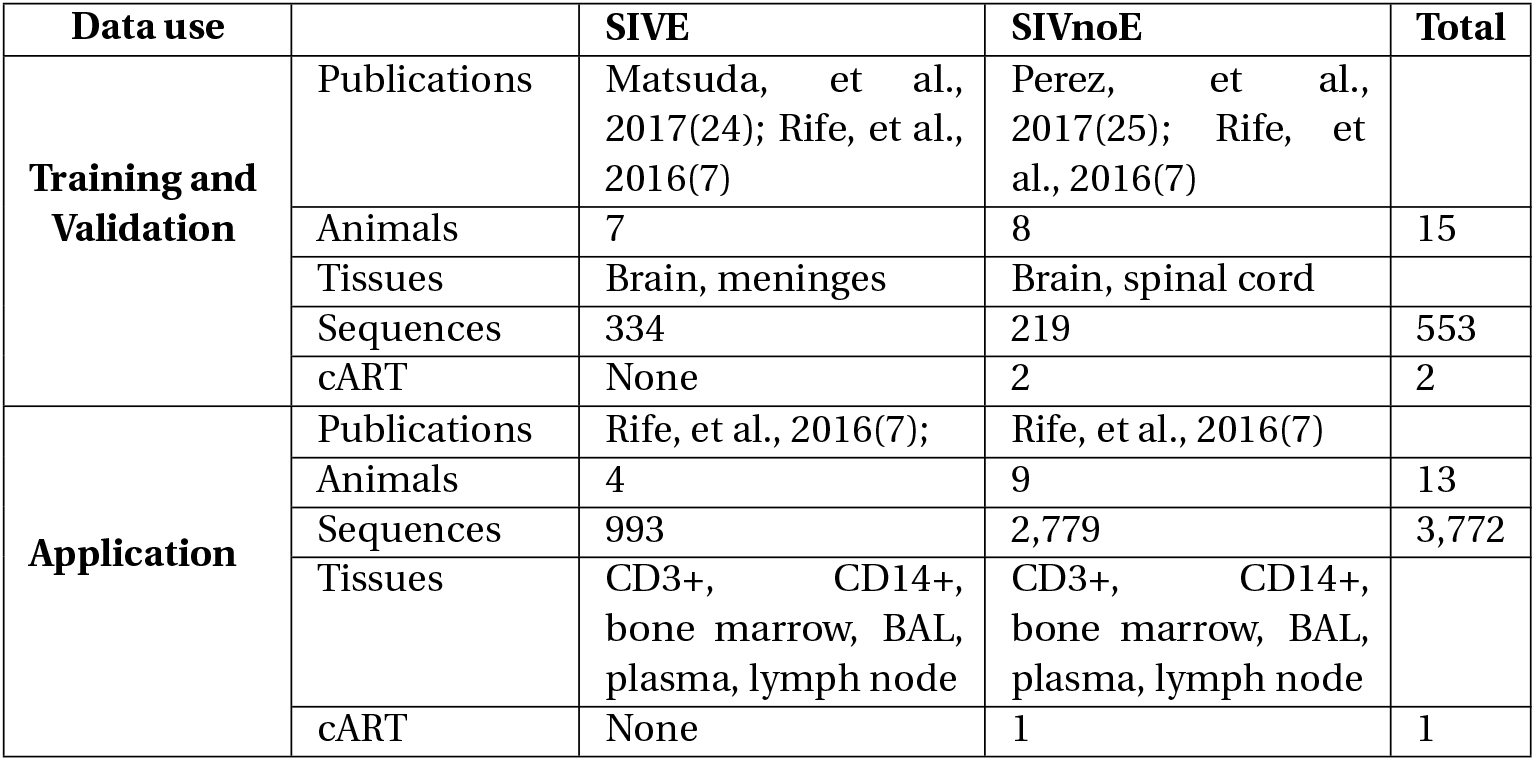
Summary of macaques included in the training, validation and application datasets.

| Data use |  | SIVE | SIVnoE | Total |
| --- | --- | --- | --- | --- |
| **Training and Validation** | Publications | Matsuda, et al., 2017(24); Rife, et al., 2016(7) | Perez, et al., 2017(25); Rife, et al., 2016(7) |  |
|  | Animals | 7 | 8 | 15 |
|  | Tissues | Brain, meninges | Brain, spinal cord |  |
|  | Sequences | 334 | 219 | 553 |
|  | cART | None | 2 | 2 |
| **Application** | Publications | Rife, et al., 2016(7); | Rife, et al., 2016(7) |  |
|  | Animals | 4 | 9 | 13 |
|  | Sequences | 993 | 2,779 | 3,772 |
|  | Tissues | CD3+, CD14+, bone marrow, BAL, plasma, lymph node | CD3+, CD14+, bone marrow, BAL, plasma, lymph node |  |
|  | cART | None | 1 | 1 |

### Machine learning to extract genetic signatures

We applied a ML approach based on Holman and Gabuzda (15) to SIV *gp120* sequences obtained from the CNS of animals with or without SIV-associated encephalitis. We proceeded according to the following three main steps: (1) feature/attribute selection (independent for each fold in cross-validation), (2) classification and model selection, and (3) prediction assessment, wherein steps 1 and 2 were used in model training, and step 3 served as evaluation of the predictive performance (26). A schematic summary of the implemented pipeline is illustrated in Figure 1. A detailed description of the approach can also be found in the Materials and Methods. Scripts used in the training, validation, and application of the model are available in https://github.com/salemilab/neuroSIVirulence.

**Fig 1.**
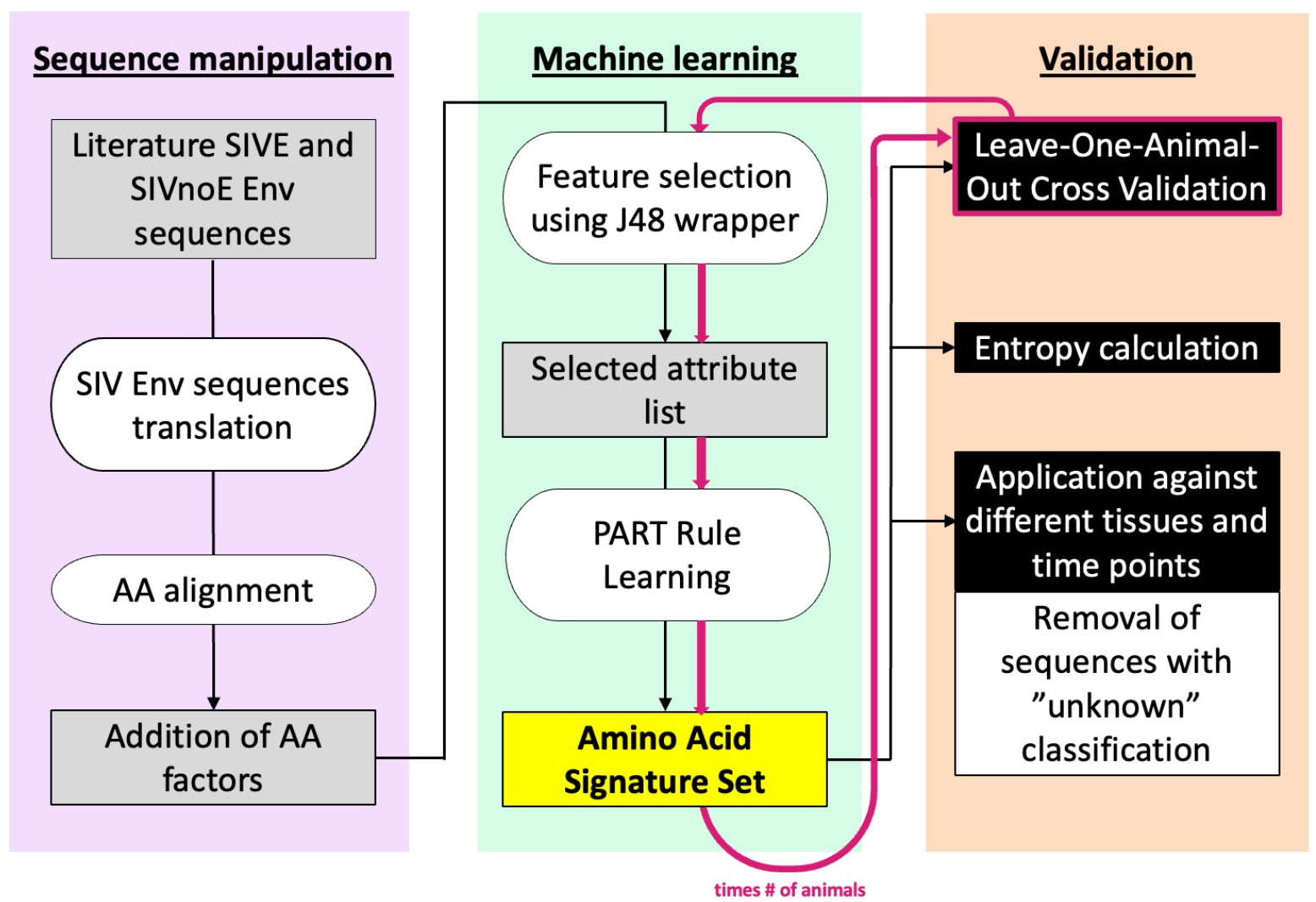
Schematic of the machine learning (ML) pipeline used in the identification and validation of viral amino acid signatures associated with SIVE. The first step corresponds to sequence manipulation wherein DNA sequences are translated and aligned, followed by annotation of each amino acid according to four biochemical features – molecular size, electrostatic charge, polarity, and secondary structure. Feature selection is then performed for selection of the most informative attributes. The final ML model is obtained from these selected attributes, and an amino acid signature set (or “rules”) is obtained. To evaluate the model, a leave-one-animal-out cross-validation was performed. The final model was then applied to sequences from different tissues and time points. Finally, we verified that signatures the model extracted are not biased towards high-entropy positions through the HIV LANL Shannon Entropy-One tool.

### Macaques as independent samples

Phylogenetic tree reconstruction was performed for aligned SIV Gp120 amino acid sequences obtained from a variety of tissues and time points (when available) from animals diagnosed with (SIVE) or without (SIVnoE) SIVE. Phylogenetic clustering according to SIVE status or cohort was not observed (Figure 2), indicating the absence of cohort-specific selective pressure and significant convergent evolution of a neuroadaptive genotype, similar to (15). Sequences sampled from different animals at earlier time points during infection intermixed in the phylogeny, as expected given that all animals were infected by the same viral swarm (SIVmac251) (27). Conversely, evolutionary divergence at later time points resulted in macaque-specific clustering, indicating animal-specific genetic drift of the virus population and, thus, allowing for each animal to be treated independently within the model.

**Fig 2.**
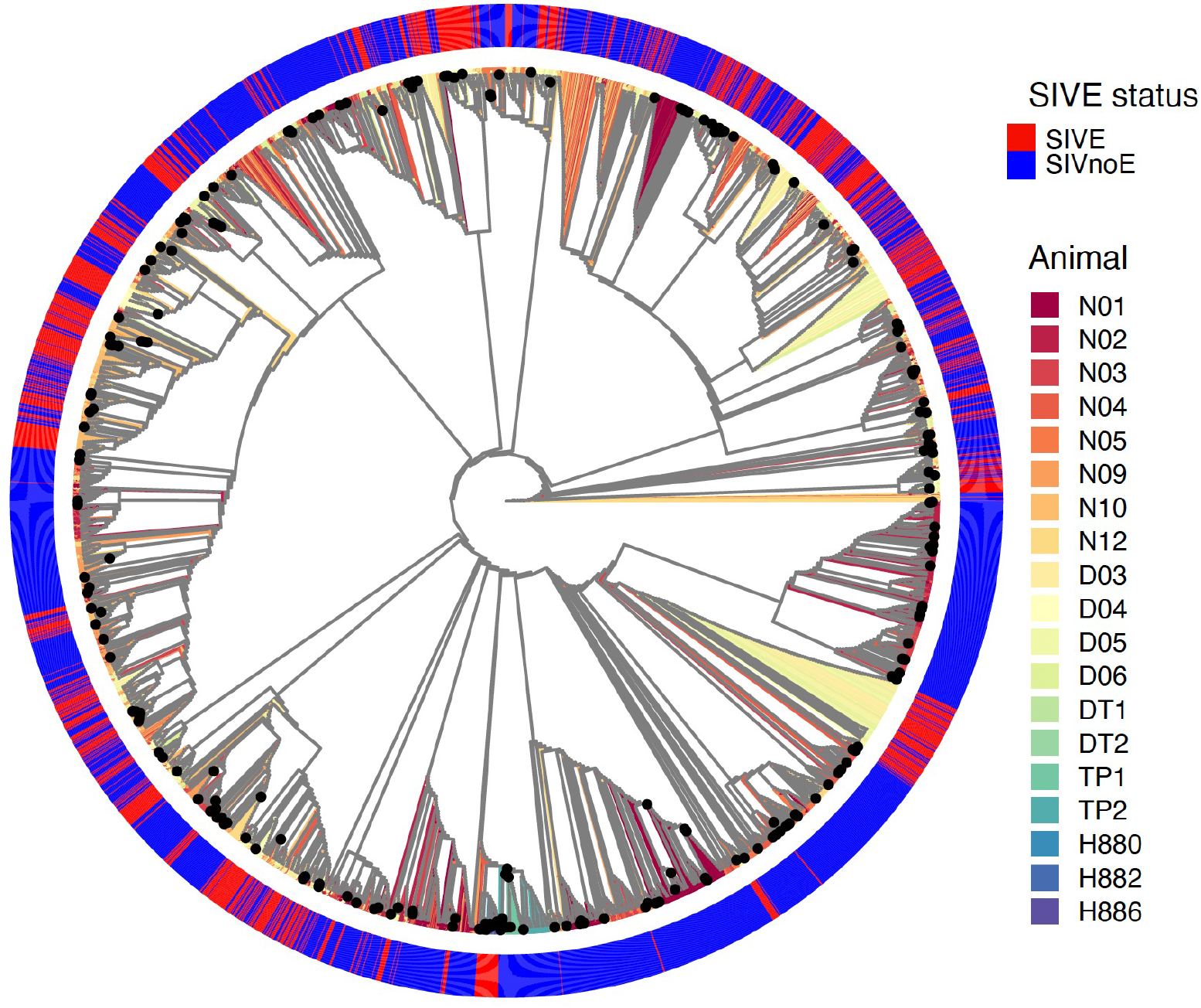
Maximum likelihood phylogenetic tree reconstructed from SIV *Env gp120* sequences from multiple tissues and time points through infection from animals diagnosed with (SIVE) or without (SIVnoE) SIV-associated encephalitis.

### Predictive genetic signatures of SIVE status and model performance

Individual amino acids within necropsy sequences originating in the CNS (including brain, meninges, and spinal cord) were assigned the following biochemical properties as described in Atchley *et al.*(28) – polarity, electrostatic charge, molecular size, and secondary structure (also used by Holman and Gabuzda (15)). The machine learning prediction model was then trained using the projective adaptive resonance theory (PART) rule-learning algorithm, resulting in a collection of amino acid signatures. The individual rules obtained within the set of rules given by the learned model were interpreted as amino acid signatures. Performance of the model were estimated using leave-one-animal-out cross-validation (Figure 1).

The final model extracted a total of nine rules, of which four were predictive of SIVE (Figure 3) and four of SIVnoE. An additional, default rule was generated for automatic classification of sequences if no other rule applied. The default rule was not considered, however, in our analysis, which is focused on extracting signatures from the rules. Thus, sequences not classified as SIVE were considered to fall under the category of SIVnoE. SIVE rules were comprised of a combination of different amino acid positions, as well as an assortment of amino acid identities and their biochemical properties considered for the model. Following feature selection, amino acid identity, polarity, molecular size, and role in protein secondary structure, but not electrostatic charge, were considered the most informative for SIVE prediction. This finding was in contrast with that of Holman and Gabuzda ((15)), who reported all four biochemical properties as important features. Most of the signatures involved amino acid positions within the variable regions V1 to V4 of SIV Gp120. When applied to the validation data set per monkey, these rules demonstrated 75% precision and 83% balanced accuracy (Table 2), with signature 1_01 having classified (as SIVE) the greatest number of true SIVE animals (4) and CNS sequences from SIVE animals (193) (Table 3). Additionally, recall, true negative rate (TNR), and F1 score were all above 70%, indicating an adequate model for SIVE prediction.

**Fig 3.**
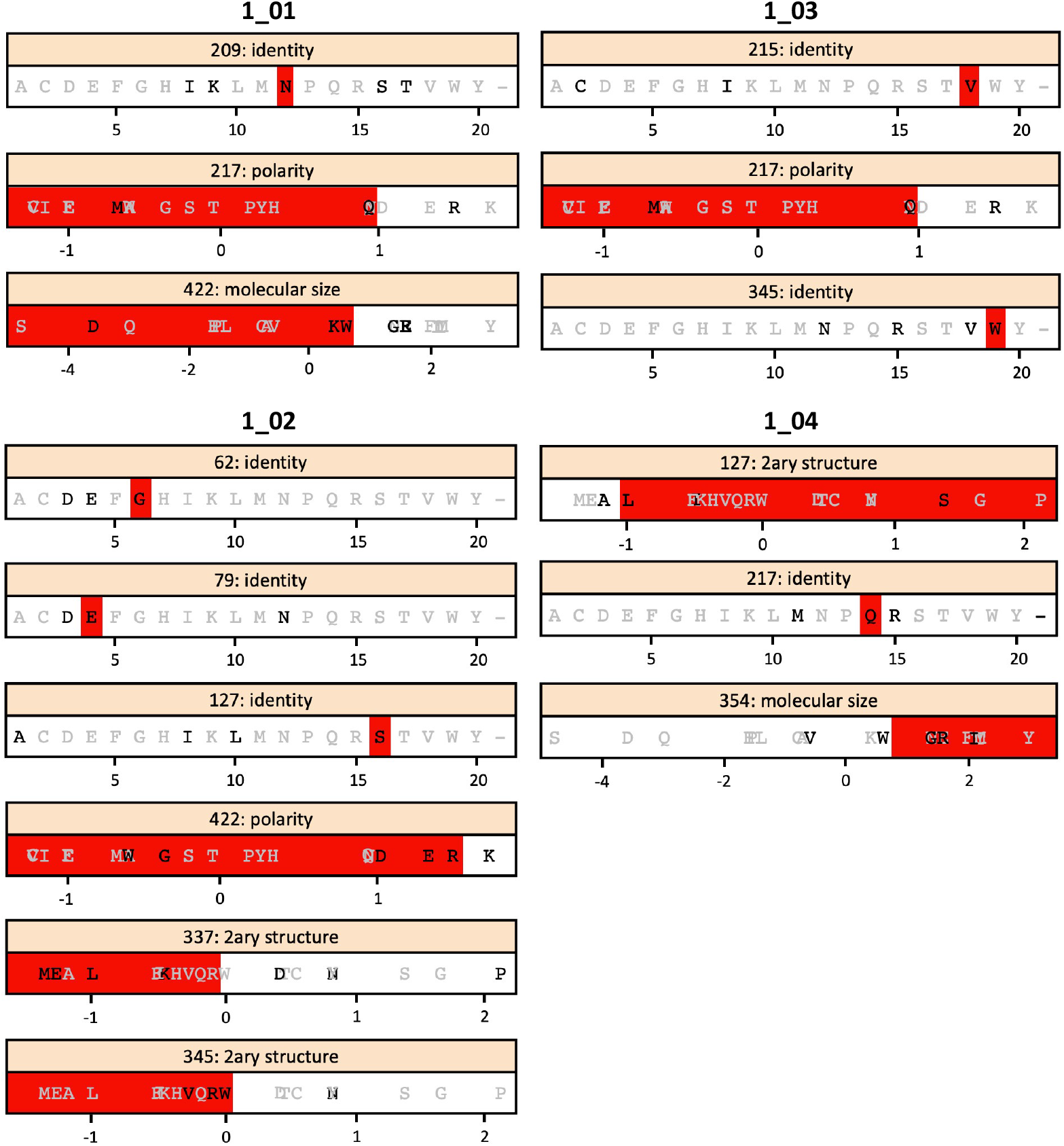
SIVE rules based on the machine learning model used for necropsy-sampled CNS sequences from SIVE and SIVnoE animals. For each position, amino acids observed at that position are colored in black, positions not observed in gray. The red bar indicates the range of acceptable values in that signature, according to the obtained rules using PART algorithm. Positions given are based on the SIVmac251 accession KU892415.1 envelope protein, id AMX21539.1.

**Table 2.** Performance statistics of the machine learning model generated with the validation data set consisting of necropsy-sampled CNS sequences from SIVE and SIVnoE animals.

| Statistic | Validation set per sequence | Validation set per animal* |
| --- | --- | --- |
| Precision | 0.82 | 0.75 |
| Recall | 0.71 | 0.86 |
| True negative rate | 0.76 | 0.8 |
| Balanced accuracy | 0.76 | 0.83 |
| F1 score | 0.74 | 0.8 |

**Table 3.** SIVE classification in animals diagnosed with SIVE.

| Signature | Number of sequences | Number of animals |
| --- | --- | --- |
| 1_01 | 193 | 4 |
| 1_02 | 60 | 3 |
| 1_03 | 26 | 4 |
| 1_04 | 3 | 2 |

Amino acid site 217 was the most frequently observed (three of the four rules, including 1_01), though feature importance varied between identity (glutamine allowed) and polarity (<1). This amino acid is located between variable regions V2 and V3 of the Env protein (Figure 4A). Similar findings have been reported previously (7), whereby an increase in the number of sites under selective pressure was observed within the V1 and V2 regions of sequences from SIVE animals, though increased selection was also observed within the constant region C2 (7). The V2-V3 region also contained the greatest number of identified sites (3), suggesting an important role for this region in neuroadaptation. In general, however, the four rules were not restricted to one particular region within the protein, though V5 was excluded in all rules. Results resembled those of the study by Holman and Gabuzda (15), which, although limited to the C2-C3 region, uncovered similarly consistent overlap of V3 sites among signatures. The increased presence of V3 sites among signatures in this previous study could not be explained entirely by mutational bias, as positions included in signatures exhibited a wide range of mutational entropy (15). Consistent with this finding, we observed a similar wide range in entropy values across signature sites (Figure 4B), suggesting sites experiencing high evolutionary rates were not restricted to neuroadaptive function and that the model was not biased towards mutable sites. While signature sites experienced approximately 2.5-fold higher entropy on average than sites not designated as signature sites, this difference was not significant (p<0.1) and is likely attributable to the difference in number of positions evaluated in each, with positions in signatures being considerably lower in number as compared with the positions not observed in signatures (10 positions in signatures versus 442 positions not in signatures).

**Fig 4.**
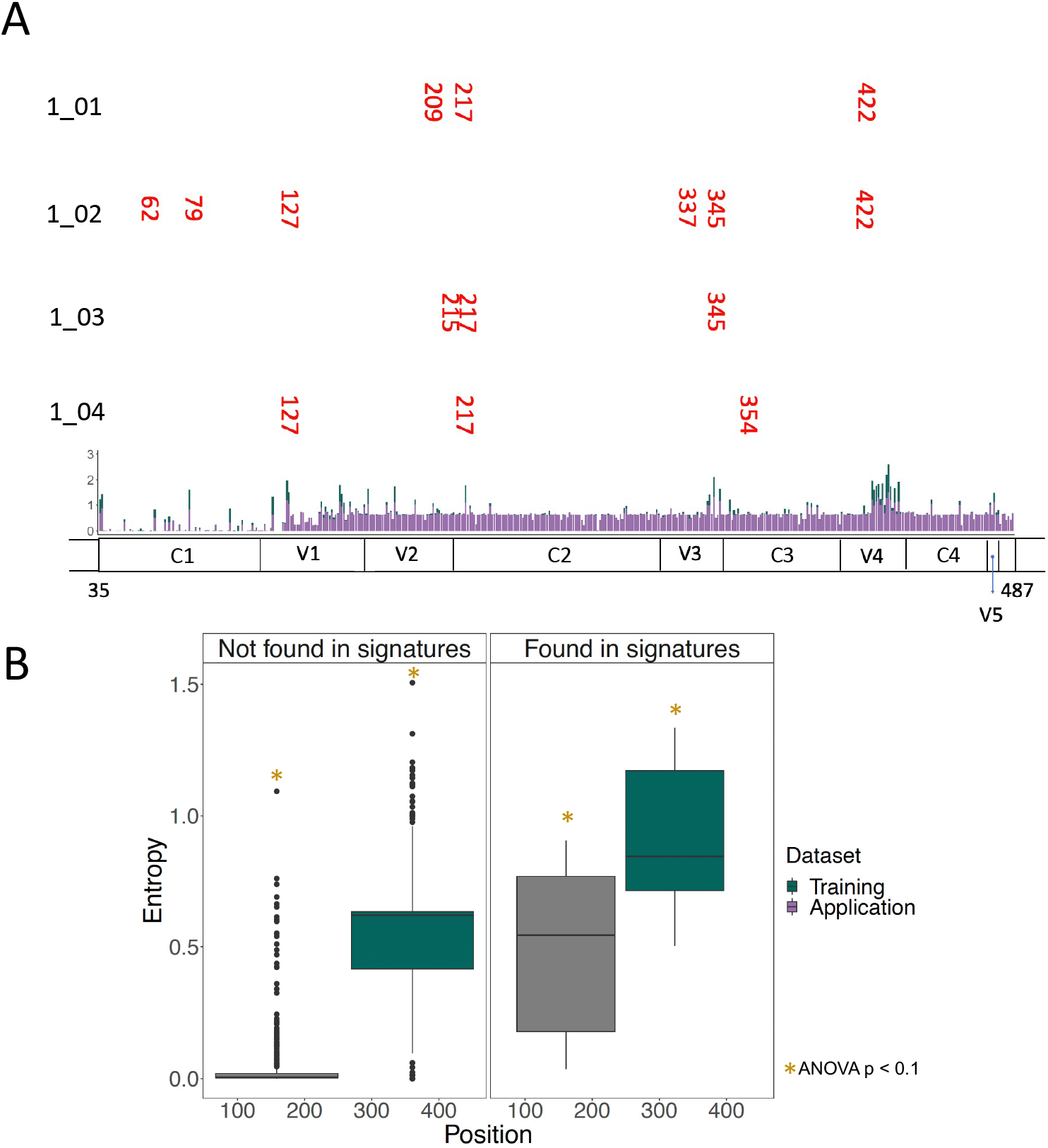
Amino acid positions and corresponding calculated entropy. (A) Shannon entropy values for each amino acid in the training (green) and testing (purple) datasets. Signature-associated amino acid positions are also highlighted in red text above their respective genomic location (x-axis) and adjacent their corresponding signature(s) (y-axis). SIV Env constant and variable regions are also shown along the x-axis. (B) Entropy value distributions among amino acid positions according to SIVE signature association.

In order to evaluate further the potential mutational bias of the PART algorithm, we performed a Bayesian graphical model (BGM) analysis of pairwise mutational associations (i.e., co-evolution of sites) between amino acid acid positions. This approach was specifically used to determine if the tendency for higher entropy sites to be included in SIVE signatures could be explained by epistasis, represented by frequent co-evolution of sites along the same branch within the phylogeny and often observed during adapting to new environments (29). Posterior probability of at least 90% for site pairs within the model was considered supportive of co-evolution (Figure 5). Seventy-nine total sites were considered co-evolving, including a minority of four signature-associated sites; however, none of these sites occurred as co-evolving pairs, indicating similar mutational patterns across sites were not responsible for PART classification. While co-evolution of two or more signature sites within the same rule might be expected to preserve neuroadaptive function, observance of co-evolution of signature sites with non-signature sites is not entirely unexpected – amino acid residues involved in neuroadaptation likely perform other functions at the protein level (e.g., proteir folding) on which other non-signature residues may be conditionally dependent.

**Fig 5.**
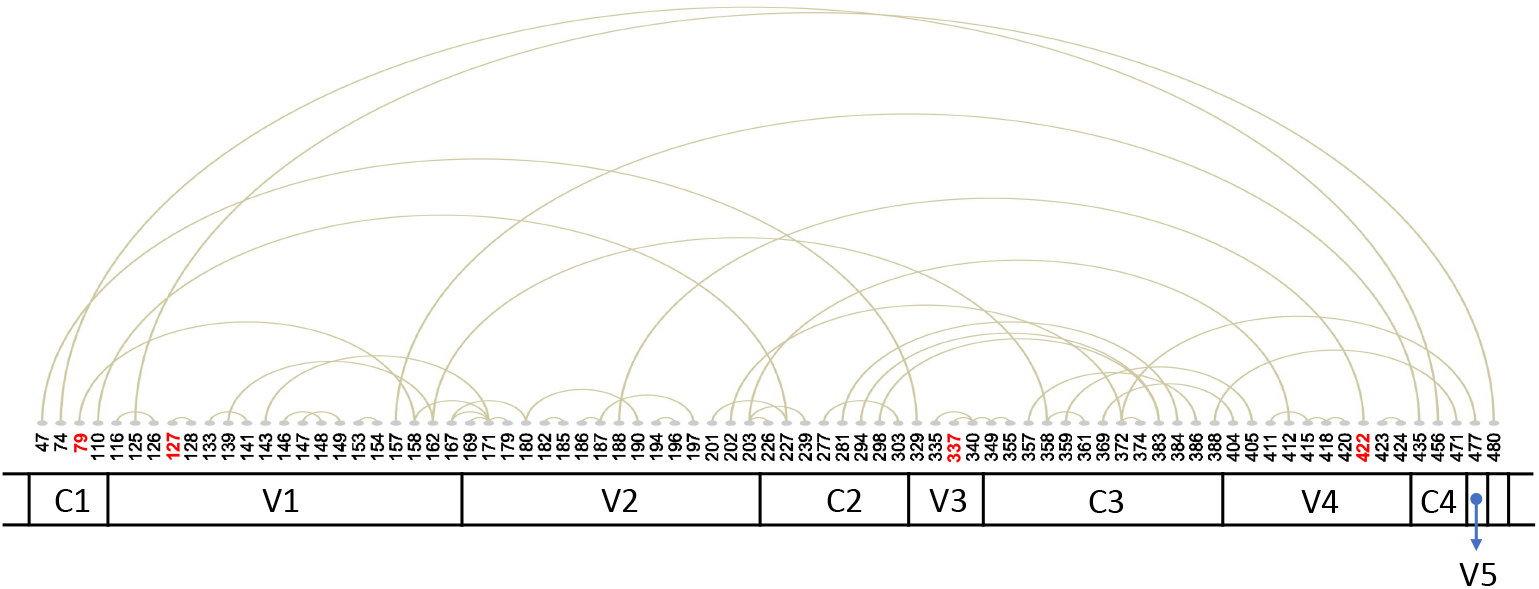
Graphical representation of conditional dependencies obtained from the Bayesian graphical model for SIV Gp120 amino acid sequence mutations. Arcs represent the association of two SIV Gp120 amino acid positions exhibiting co-variation. Diagram on the bottom shows the SIV Gp120 constant (C) and variable (V) regions. Position numbers based on SIVmac251 Gp120 reference sequence (accession KU892415.1, protein id AMX21539.1). Only probabilities ≥90% are shown. Positions in red are positions found in SIVE signatures.

### Spatiotemporal distribution of SIVE-associated amino acid signatures

The learned model was next applied to sequence data from additional sampled tissues and time points in order to gain better insight into how early viral-mediated neuropathology can actually be predicted and the invasiveness of the sampling procedure required for reliable SIVE prediction. All remaining tissues (listed in Table 1) and time points for animals described in Rife *et al.*(7) were used in this application. The observance of SIVE signatures in tissues outside the CNS and at earlier time points, at times as high as 78% of sequences (Figure 6) was initially promising, as it demonstrated the potential for prediction of neuropathology without invasive collection of CNS samples and potentially at a time point earlier than pathological onset that, if early enough, may provide sufficient time for therapy-mediated prevention. However, SIVnoE animals also exhibited the presence of SIVE signatures at earlier time points. A high rate of SIVE classification among early sampled sequences in both cohorts, particularly for signature 1_0_2, might not necessarily be a misclassification of the model, but rather an indication of the presence of SIVE signatures in the infecting viral swarm (SIVmac251 ((30))), which was originally obtained from monkeys at the time of necropsy. It is possible that this infecting population/quasispecies may actually harbor SIVE signatures that are not beneficial at the time of, or immediately following, infection/transmission.

**Fig 6.**
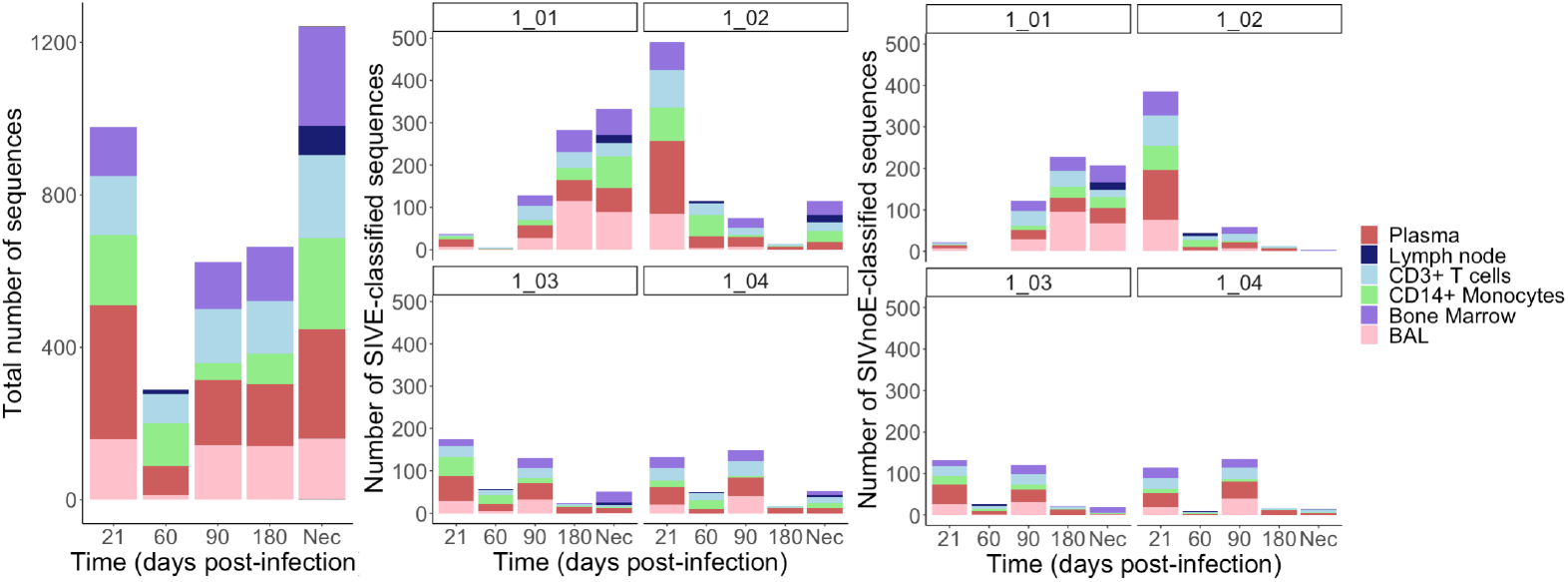
Classification of sequences obtained from several tissues at different time points in each group (SIVE or SIVnoE) of animals. (Left) Total number of sequences collected from each tissue at each time point. (Center) Number of sequences classified as SIVE from SIVE animals for each tissue at each time point split by SIVE signature. (Right) Number of sequences classified as SIVE from SIVnoE animals for each tissue at each time point split by SIVnoE signature.

A high degree of presence of SIVE signatures in non-CNS tissues was not anticipated, considering CNS sequences have historically been phylogenetically distinct from sequences collected outside the CNS (8; 31; 32; 33). The presence of shared genetic signatures across these otherwise compartmentalized sequences indicates that, in the face of natural evolution of the virus in the CNS, some specific protein property required for CNS entry and/or replication is not entirely unique to the CNS. The ability of the virus to adapt to new cell types, for example, is well known. The development of macrophage tropism is particularly well-characterized and has been linked previously to neuropathology (34; 35). As the primary cell types infected within the CNS are perivascular macrophages and microglia (6), it is not unreasonable to rationalize that cellular tropism is a necessary phenotype for CNS infection. Indeed, lung macrophages comprising the bronchoalveolar lavage fluid (BAL) were the only tissue population to classify 100% of sequences as SIVE in SIVE-diagnosed animals using this machine learning model. As this population was also classified as SIVE at high levels in SIVnoE animals at earlier time points, we posit that macrophage tropism is not the only requirement for neuroadaptation. Moreover, macrophage tropism has been reported in a variety of individuals, regardless of neuropathology (36), suggesting that macrophage-tropic variants might be established and replicating in macrophages before neurological pathology becomes apparent.

### Structural similarities between SIV and HIV Gp120 signatures

Next, we wanted to examine the extent to which the results of Holman and Gabuzda (15) in PLWH were translatable to the SIV-macaque model. Similar to HIV, SIV makes use of envelope glycoproteins to bind to cellular surface receptors and infect their target cells. Although macaque SIV and HIV-1 share only ~ 35% sequence identity in their envelope glycoproteins, they exhibit higher similarity in other, more functional aspects (e.g., 70% similarity in the location of disulfide bonds), and their molecular architectures are highly comparable (37). Hence, direct translation of amino acid position, or even biochemical property within the neighboring area between viruses was not anticipated. Instead, we used published Env Gp120 protein structures for HIV-1 (PDB 2NY3 ((38))) and SIV (PDB 3JCC ((39))), highlighting the signature-associated amino acid positions obtained from Holman and Gabuzda and herein, respectively in order to look for structural clues in terms of functional similarity (Figure 7). Overall, signatures for both HIV ((15)) and SIV could be categorized as exposed within their respective molecular structures (Figure 7), with 1_04 being partially exposed in SIV In HIV 80% of the signatures determined by Holman and Gabuzda ((15)) were involved in contacts with NAG. The HIV PDB structure held contextual information regarding 2-acetamido-2-deoxy-beta-D-glucopyranose (NAG) residues, revealing information regarding putative aminoglycan contacts among amino acid residues. A distance of 4.5Å between amino acid and NAG was considered evidence of putative contact dependency. When mapped, HIV-1 demonstrated five signature-associated amino acid residues with putative NAG contacts (Table S1), all within the C2 region of Env. For our SIV sequences, position 345 (332 according to PDB numbering) was identified in two of our signatures: requiring the amino acid tryptophan (W) per SIVE rule 1_03 or a secondary structure equal or below 0.009 (according to the AA index values, Figure 3) per rule 1_02. This amino acid position, representing the V3 region boundary, was considered within contact distance of an NAG. Additional two amino acid positions (e.g., 97 and 401) from our signatures were considered to be in relatively close proximity to NAG, but not within the considered 4.5Å distance. It is important to note that SIV NAG contacts were inferred from the HIV PDF (i.e., based on the superimposition of the HIV and SIV Gp120 structures), as the resolution for the cryo-electron microscopy SIV Gp120 structure was insufficient for NAG placement. Subtle differences in the HIV and SIV Gp120 structures (Figure 7) may translate to differences in NAG placement, resulting in an underestimation of the number of true NAG contacts. As an example, residue 401 in SIV pattern 1_0_1, position 401 is located within a region that does not align well with HIV structurally, yet this residue is still in close proximity to the corresponding NAG. Among the SIV rules, 1_0_4 was the only signature set of residues not clearly interacting with NAG molecules, which accounted for the fewest SIVE-classified animals and sequences during model training.

**Fig 7.**
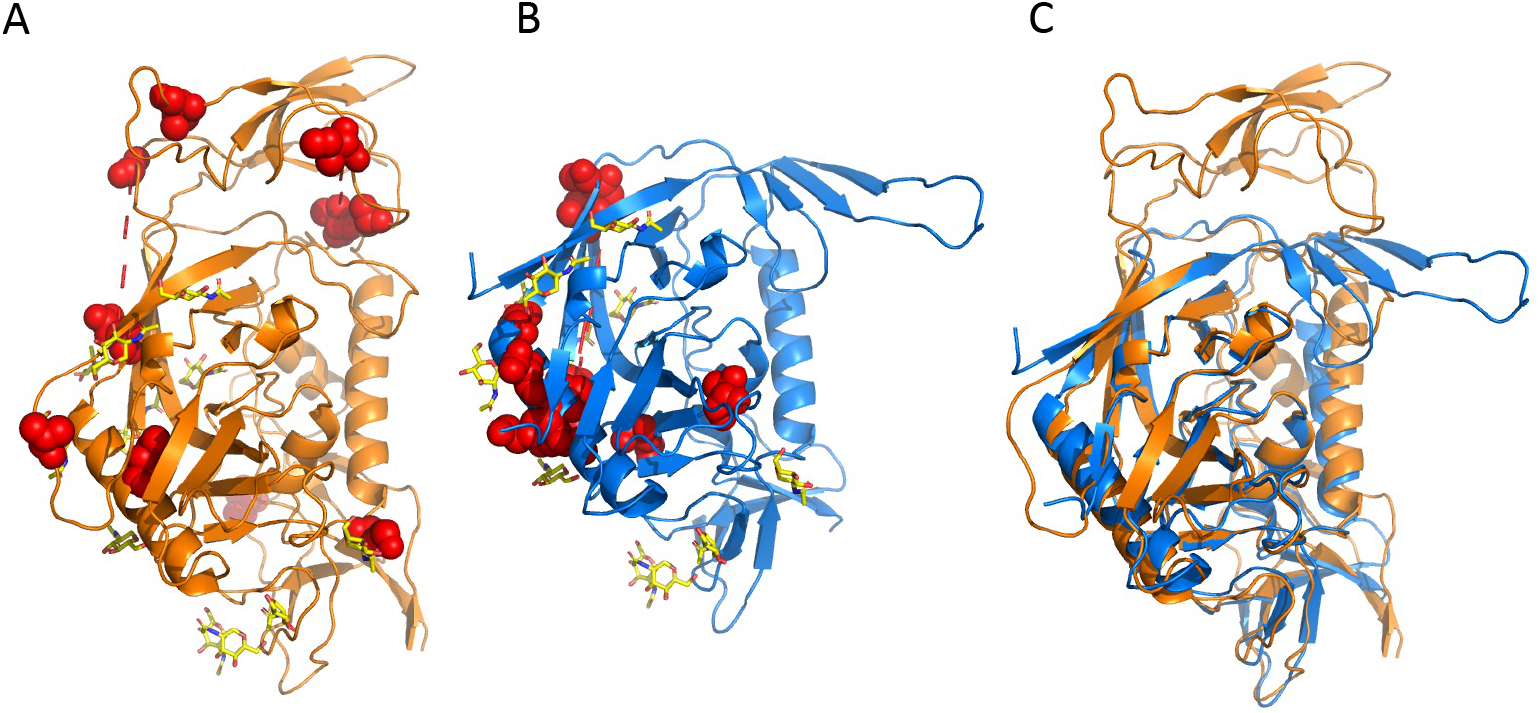
Amino acid signatures mapped to HIV and SIV Env gp120 3-dimensional protein structures. (A) SIV envelope crystal structure (PDB 3JCC). (B) HIV envelope crystal structure (PDB 2NY3). (C) Overlap of SIV and HIV envelope crystal structures from (A) and (B). Red spheres represent positions found in SIV and HIV amino acid signatures.

### Phyloanatomic inference of the source of neurovirulent virus in the brain

While a shared macrophage-tropic phenotype between virus in the CNS and lungs is the simplest explanation for SIVE classification of both tissues, another explanation is that the source of SIVE signatures in the brain is the actual migration of infected macrophages from the lungs to the CNS. Epidemiological modeling within a phylogenetic framework within a host, referred to as phyloanatomy (20), offers an *in silico* solution to understanding both viral evolution and dissemination of virus among various anatomical compartments. Derived from the similar framework used to study regional and global migration of pathogens during an epidemic, phyloanatomic analysis assumes a network of isolated tissues tied together via the vascular system and circulating immune cells carrying the virus. Using serial sampling during the course of infection, and assuming a clock-like model of evolution, this method is also able to resolve the timing of relevant evolutionary and epidemiological events, such as the question of whether early viral entry followed by isolated replication in the CNS or late entry of a neuroadapted viral variant that emerged in the periphery is responsible for neurovirulence.

Pathological evaluation of HAND in humans is only possible by *post mortem* analysis of brain tissues, which provides a snap-shot of end-stage pathology, while mechanisms that contribute to onset and progression of this pathology remain elusive (40). Less than 2% of the total body lymphocyte population resides in the peripheral blood, thus highlighting the role of viral infection in deeper tissues and their potential role in seeding the brain, yet, serial sampling from all possible tissue sources of HIV CNS infection in humans is an additional hurdle when not a practical impossibility. Sampling from tissues at multiple, pre-determined time points is more readily available for animal models, which have provided valuable insight into the plasticity of tissue infection (41). Previous SIV sequence analyses performed by our group have indicated a potential role for multiple tissues in seeding the brain (27) and a prominent role for non-CNS tissues in the evolution of a neuroadapted virus that enters the brain during the late stage of infection (7). The specific tissue(s) responsible for entry of virus into the brain during early and end-stage disease have not yet been identified in the context of neuropathology but are readily discernable using statistical phyloanatomy techniques (42; 43). Complementing the identification of amino acid signatures of virus capable of replicating in the brain, inference of the viral dissemination patterns among the plethora of infected compartments within the encephalitic host is a critical step toward predicting the risk for development of cognitive impairment. By employing the Bayesian phyloanatomy framework(20), we investigated potential source(s) of CNS virus in the two animal cohorts described in the machine learning application above, revealing insight into differences in the source and timing of entry of virus between animals that develop SIVE and those that do not.

Bayesian analysis of viral diffusion over the course of infection in SIVE and SIVnoE animals revealed that the virus may be more readily accessible to certain tissues depending on the duration of infection. In both animal groups, viral dissemination during the early phase of infection (first 21 dpi) was primarily driven by exchange between monocytes, T cells, and blood, as well as from bone marrow to the peripheral T-cells, with few exceptions (Figure 8). One notable exception was the early circulation of virus among the individual brain cortices, or lobes, of the SIVE animals, suggesting earlier brain infection in these animals than in animals without SIVE (SIVnoE). As with all dispersion pathways depicted in Figure 7 and discussed hereafter, this circulation was considered significant *(BF >* 3), characterized by flow of virus from the parietal to the frontal cortex, although the significant anatomical source could not be identified. Following the period of early infection, viral dispersion patterns in the periphery diverged, differing from early infection for both animal groups. During the designated asymptomatic time interval, SIVnoE animal sequences exhibited limited dissemination between, as well as to, PBMCs and plasma. Significant exchange between PBMCs and plasma continued into asymptomatic infection, however, for SIVE animals, with the additional contribution of virus from lung macrophages. Viral dissemination among the three parenchymal cortices and meninges was detected during asymptomatic infection for both groups, with no significant identifiable anatomical origin.

**Fig 8.**
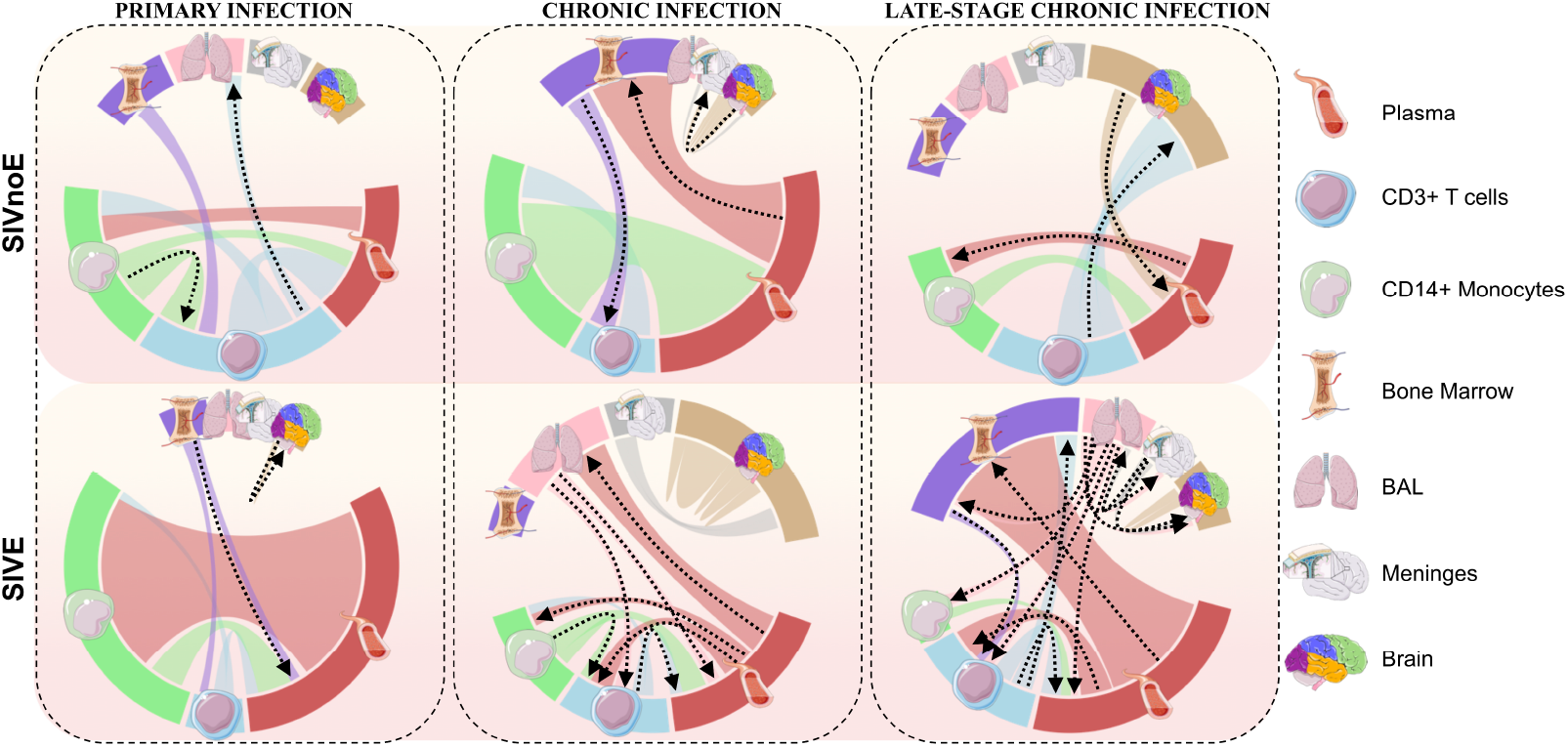
Graphical representation of viral dispersion patterns over the course of infection in the context of SIVE for the SIV-infected macaque model of HIV infection and neuroAIDS. Bayesian stochastic search variable selection (BSSVS), assuming asymmetric diffusion among discrete anatomical compartments, was implemented in BEAST v1.8.3. The hierarchical phylogenetic model was used to infer spatiotemporal trends in dispersion across animals with (n=4) and without (n=3) SIVE. Diffusion model parameters were allowed to differ between designated time intervals corresponding to early infection (21 days post-infection), AIDS onset (21 days prior to necropsy), and asymptomatic infection (time span between early infection and AIDS onset). Arrows indicate directionality of significant (Bayes Factor >3) diffusion and are colored according to tissue of origin.

AIDS-related neuropathogenesis in SIV-infected macaques is characterized by distinct viral dispersion patterns during the late stage of infection. As dispersion patterns continued to diverge from the early infection period and between the SIVE and SIVnoE macaque groups, the onset of AIDS was marked by notable differences in not only the origin of brain viral sequences but of connectivity between peripheral tissues. A loss of significant exchange of virus between peripheral monocytes and T-cells was detected for both groups during this time interval as compared with earlier infection. Also in contrast to earlier time intervals, meninges contributed significantly back to the periphery in both animal groups, specifically the peripheral blood and lungs in the SIVnoE and SIVE animals, respectively. It is important to note that as brain and meninges sequences were only available upon necropsy, the exchange of cranial and meningeal viral populations with those of peripheral tissues may be underestimated for earlier time points. However, nearly uniform sampling over time for the remaining tissues and cell populations offers greater confidence in the indication of a highly dynamic viral population network over the course of SIV infection.

Despite similarly limited contribution of peripheral tissues and cell types to infection of the T-cell population, an increased number of dispersion pathways originating from peripheral T-cells was observed for the SIVnoE animals as compared to early infection. Alternatively, for the SIVE animals, a larger transmission network was observed compared to earlier time periods. These transitions consisted largely of movement into the bone marrow, PBMCs, and peripheral blood, almost exclusively from the lung macrophages. This family of macrophages was also the exclusive contributor among sampled locations to cranial virus in SIVE animals, with this exclusion reserved for peripheral T-cells in SIVnoE animals, both of which were only significant during late infection.

## Discussion

In this study, we expanded existing machine learning methods(15; 19) for application to the animal model of HIV-associated neuropathology in order to identify genetic signatures in the envelope (Env) Gp120 protein correlated with the onset of neuropathology. We identified four amino acid signatures associated with the presence of SIVE from CNS sequences obtained at necropsy and utilized these signatures to predict SIVE relevance for sequences amplified from plasma, lymph node, bone marrow, bronchoalveolar lavage fluid (BAL), and peripheral blood monocytes and T cells, each sampled at different time points during the course of infection and/or treatment. Our goals were to 1) assess the translational significance of such signatures in PLWH to the macaque model, 2) relate common signatures to protein structure and function, and 3) to determine if neurovirulent viral variants could be identified outside the spatiotemporal domain of the CNS at the time of symptom onset for more clinically relevant prediction of risk.

Amino acid residues identified in signatures predictive of SIVE using our model were not localized to any sub-region of the Gp120 protein, consistent with previous findings (15), with the limitation of only evaluating the C2V3C3 region (19), and acting to explain the seemingly contradictory findings of numerous studies attempting to identify minimal signatures of neurovirulence (12;44;45;46). Regardless of non-localized signals, residues within the identified SIVE signatures tended to occur within hypervariable regions (V1, V2, V3 and V4). Dehghani *et al.* (2003) observed genetic variation during the course of infection in rapidly progressing macaques within these regions associated with the distinction of brain virus from peripheral blood, representing one of the earlier studies presenting evidence in favor of a potential role for viral evolution in neurological disease progression (47). Dehghani *et al.*specifically noted increased variability for V3 residues 340 and 348 (337 and 345 in SIVmac251 reference and observed among our signatures), proposing affected binding affinity of Gp120 for the CD4 cellular receptor. Differential binding of this receptor is required for more effective replication within cells with lower abundance of surface CD4 molecules, such as monocytes/macrophages (48), which comprise the majority of the infected population within brain tissue (49). Indeed, replacement of valine with aspartic and glutamic acid at position 337 was observed in 100% of SIV sequences extracted during late-stage infection in brain, bone marrow, and lymph node from our *Rhesus* macaques. Though valine was not one of the amino acids observed at this position among our sequences, it is in compliance with our SIVE rule for this position regarding secondary structure. These findings corroborate the validity of our model and its translation in HIV infection, encouraging increased efforts to expand on the repertoire of residues involved and/or interacting with this region (V3) in the development of neurovirulence.

Ogishi Yotsuyanagi (2018) similarly expanded on the Holman and Gabuzda study, incorporating a more sophisticated validation approach in their machine learning model and more comprehensive dataset. While Ogishi Yotsuyanagi applied a hold-out validation in which they sought to correct the leave-one-sequence-out generalization error from the Holman and Gabuzda study, our cross-validation was explicitly designed to assess the model capabilities to generalize. By using a test fold constituted by all the sequences from a single animal, we took a conservative stance, as virus populations from the same animal evolve together and, therefore, cannot be considered independent. With our design, we avoided the risk of our estimates to be as optimistic as they could be, for example, with a leave-one-(sequence)-out or traditional cross-validation that does not consider the sequence origin (i.e., sequences from the same animal are present in both test and training sets). In an attempt to increase the robustness of the model, Ogishi and Yotsuyanagi used all sequences available from each diagnosed individual within the Los Alamos National HIV Database (50), including blood, lymphatic tissue, and others. As demonstrated in our study, however, disease signatures represent a fraction of the total number of sequences in an individual and may be present in other tissues. Moreover, several studies have demonstrated compartmentalization of sequences in non-CNS tissues (36; 51; 52;53;54;55) or even cell types (56; 57), suggesting the introduction of a significant amount of noise with the inclusion of multiple tissues and cell types.

Capable of transmigration across the blood-brain barrier (BBB) and susceptible to viral infection, mature monocytes are believed to play a key role in facilitating the chain of events that result in neurocognitive impairment (49). The expansion of the mature (CD14+CD16+) monocyte subpopulation from 5-10%of all peripheral blood monocytes to ~ 40% in HIV-infected individuals is predictive of HIV-associated neurocognitive decline and may increase the likelihood of these events (58; 59). While monocytes may act as Trojan horses in the delivery of virus, they may not act alone. The high prevalence of SIVE signatures observed in this study in lung macrophages, and statistical phyloanatomic evidence of these cells as ancestors of viral progeny in the brain, suggests a supplementary source of brain infection while still consistent with previously described association of neuropathology with a macrophage-tropic phenotype. Infecting viral strain SIVmac239 was reported to replicate poorly in primary alveolar macrophages derived from lung lavage samples of healthy virus-negative monkeys, but extremely well at necropsy (168 dpi), indicating this phenotype evolves over the course of infection (60). Similarly, whereas few BAL sequences were classified as SIVE during initial infection in this study, sequences classified as SIVE according to the most prominent signature (1_0_1) grew in number over time. The presence of these signatures as early as 21 days post-infection, during which the viral population is difficult to discern genetically from the original infecting swarm, suggests that minor variants within the original infecting population may be capable of causing disease but are less fit than the remaining population. Following the emergence of this phenotype in deep tissues such as the lungs, recruitment of pro-inflammatory macrophages and lymphatic connection to the CNS may result in increased flux of virus.

HIV isolates capable of efficient replication in the brain, deemed “neurotropic”, exhibit increased sensitivity to neutralizing antibodies, revealing the trade-off between cellular tropism and immune evasion (11) and providing an explanation for the low fitness of this phenotype during early infection. Hypervariable regions V1, V2, and V3 have been implicated in this trade-off (61;62), consistent with our SIVE signature locations. Whereas identity, polarity, and molecular size were demonstrated to be important features at these locations (particularly for signature 1_0_1), a common denominator when taking three-dimensional structure into account across HIV and SIV signatures of disease were interactions with glycan residues. Asparagine(N)-linked glycosylation of HIV Gp120 has been proposed as a significant mechanism for minimizing the virus immune recognition (63; 64; 65), but glycosylated sites have also presented patterns suggestive of selective advantages for replication in the brain, particularly in the V1 and V4 regions (54). Brese and Gonzalez (2018)(54) reported that the number and location of N-site patterns were much more conserved in the brain than in remaining lymphoid-derived tissues, suggesting a highly selective microenvironment and functional role for glycosylation that may be distinct from immune evasion (54). A previous comparison of viruses from different time points, including approximate time of transmission, reported less frequent glycosylation during the transmission of HIV type 1 sub-types A and C, suggesting less-glycosylated strains are preferably transmitted (63).

We recognize our study has limitations, as do many others diving into diagnosis prediction using machine learning. The number of animals incorporated into this study was fairly small, and animals undergoing treatment were not included in training and validation, which may harbor differing signatures owing to treatment-mediated selection pressure. Secondly, whereas macrophage tropism has been implicated as being necessary for neurological infection resulting in disease, there may be more than one evolutionary step required for sufficient neurotropism, which cannot be distinguished in this study. Significantly lower levels of viral diversity and divergence have been observed for macaques associated with disease compared to without disease, indicating separate mechanisms for entry into the brain and efficient replication in the brain (7). Signatures not associated with disease in animals with low-level virus observed in the brain (SIVnoE) do not automatically translate to entry-related patterns – they may in fact be unrelated to both entry and replication in the brain.

## Conclusion

Overall, we showed that, given a sufficiently informative training data set, machine learning can potentially predict SIV/HIV neuropathology before onset and without invasive collection of CNS samples, thus providing sufficient time for therapy-mediated prevention. In particular, our results highlight the contribution of lung macrophages as key players in the emergence of neurovirulent strains linked to neuropathology. Additional studies will be needed for deeper understanding of entry *versus* replication in the brain and their relationship with the development of neuroAIDS.

## Supporting Information

**Table S1.**
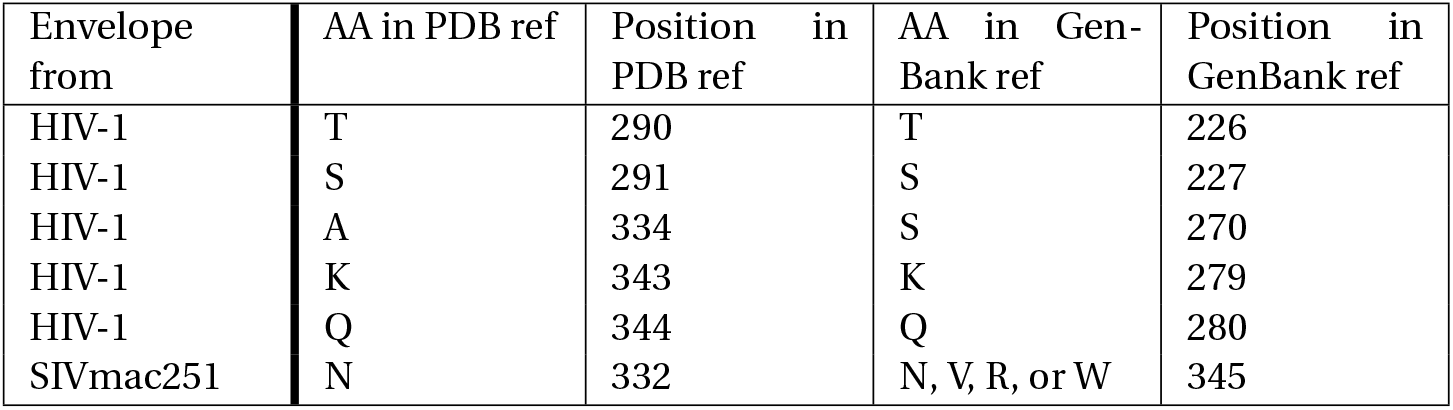
Envelope amino acid positions from HIV-1 and SIVmac251 from PDB and translation to GenBank. Genbank reference based on the HIV-1 HXB2 accession K03455.1 protein id AAB50262.1 and PDB 2NY3. For SIVmac251 accession KU892415.1 protein id AMX21539.1 and PDB 3JCC.

## Acknowledgments

The authors acknowledge the Stephany W. Holloway University Chair in AIDS Research and the University of Florida Research Computing for providing computational resources and support that have contributed to the research results reported in this publication. URL: http://researchcomputing.ufl.edu. This work was supported by National Institutes of Health and National Institute of Neurological Disease and Stroke (NINDS) grant 5R01NS063897. Andrea Ramirez was partially supported by the MacClamma fellowship.

## Materials and Methods

### Study population

Two macaque cohorts were used in this study, which included four CD8+ lymphocyte-depleted (D03-D06) and eight naturally progressing, or non-CD8+ lymphocyte-depleted (N02, N09, N10), Indian rhesus macaques (Macaca mulatta), hereafter referred to as the Mac251-DEP and Mac251-NP respectively. Both cohorts were infected intravenously with the viral swarm SIVmac251 (1 ngSIV p27) (34). CD8+lymphocyte depletion was achieved by subcutaneous administration of anti-CD8 antibody cM-T807 (6, 8, and 12 days post-infection [dpi]) (35). All animals were euthanized at the onset of SAIDS (75-300 dpi), the criteria for which included: 1) weight loss >15% body weight in 2 weeks or >30% body weight in 2 months, 2) documented opportunistic infection, 3) persistent anorexia >3 days without explicable cause, 4) severe intractable diarrhea, progressive neurological signs, or significant cardiac and/or pulmonary signs, as previously described(36). Seven of the animals harbored detectable viral sequences within analyzed frontal, parietal, and/or temporal cortex tissue sections of the brain parenchyma, as identified using single genome amplification of viral genomic RNA (described below). Pathological diagnosis of SIV encephalitis (SIVE) was determined post mortem by a veterinary pathologist and included the presence of microglial nodules and multi-nucleated giant cells confirmed by immunohistochemistry staining for SIV p27, as described previously (37–40). Animals diagnosed with SIVE included D03, D04, D05, and N10, whereas animals with detectable brain viral RNA but no detectable SIVE (referred to as SIVnoE) included D06, N02, and N09. Plasma s100B concentrations at necropsy were determined for additional SIV-infected animals with no detectable viral RNA in the brain or SIVE, which included N01, N03-N05, and N12; however, viral RNA sequences for these animals were not included in the phyloanatomy analysis, as the focus of this study was origin of brain infection. Additional macaque sequences were added, hereafter referred to as external data/sequences, obtained from published literature with clinically diagnosed rhesus macaques with SIVE(24) or SIVnoE(25) at the time of necropsy. An additional healthy macaque undergoing cART was also added.

### Nucleotide sequence accession

All of the sequences used in this study have been described previously (7; 27;30;66) and are accessible in GenBank (accession numbers JF765272-JF766081 [Mac251-DEP], KR999328-KR999727 [Mac251-NP N02 and N10], and KX081254-KX081353; KX081479-KX081498; KX081619-KX081702; KX081840-KX081862;KX082029-KX082107; KX082229-KX082252; KX082428-KX082531 [Mac251-NP N09], and MG931034-MG931480 [N03]);ON714061 – ON714132 [JA41]; SIVE external MF370654-MF370782; SIVnoE external MF284715-MF284792.

### Generation of phylogenetic tree

We aligned amino acid sequences, using Muscle in Aliview (1.26 version), from SIV-infected *Rhesus* macaques (7) diagnosed at necropsy with (SIVE) or without SIVE (SIVnoE) from naturally progressing and CD8^+^-depleted cohorts. Alignment included sequences used for model training and testing (CNS-derived), as well as sequences from additional tissues at various time points. IQ-tree (67) on command line was then used to generate a phylogenetic tree by maximum likelihood using the JTT substitution matrix.

### Weighting and translation of sequences to amino Acid properties

Following the Holman and Gabuzda method, to make sure that sequences that were obtained from different sources were weighted equally, and the number of sequence per animal was balanced, the weight per sequences was calculated by:

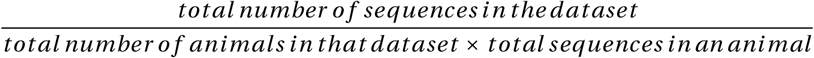

Since amino acids can have similar biochemical properties, four numeric factors (or attributes) to describe the functional role of the amino acid in each sequence were included in the analysis: polarity, secondary structure, molecular size, and electrostatic charge (28).

### Training of the model: Generation of Signatures Sets Using the PART Algorithm from Weka

We utilized sequences from rhesus macaques infected with SIVmac251 (7) as well as sequences from external data (24;25). One obstacle encountered during data set assembly when working with brain-derived sequences is sufficient sample size. We, therefore, included CNS sequences from other research groups with similarly diagnosed cohorts (24; 25). A total of 553 sequences from 15 *Rhesus* macaques (1) were used to train the model.

Feature selection from Weka (3.8.5 version) was implemented in order to use relevant features from the original set of features and to increase the performance of the model (68). We used the J48 wrapper based on information gain to extract the most relevant features (69), with WrapperSubsetEval and the BestFirst greedy hill-climbing algorithm, both with default parameters. Notably, in the cross-validation, feature selection was performed independently for each training set. After feature selection, the model was trained using the Weka projective adaptive resonance theory (PART) rule-learning algorithm with default parameters using R (4.1.1 version), to classify sequences by SIVE/SIVnoE diagnosis. PART is based on the C4.5 algorithm, a decision tree generating a hierarchical set of rules for classification(70). PART outputs an interpretable and ordered set of rules (SIVE and SIVnoE). Each rule includes amino acid requirements with which a sequence needs to comply in order to be classified. Since diagnosis is determined at necropsy, we generated the PART model using tissue sections from the brain, meninges and spinal cord taken at the necropsy time point. This model resulted in a total of nine hierarchical rules for classification. The individual rules obtained within the set of rules were interpreted as amino acid signatures. The last rule applies a default classification, if no other rule successfully matches the amino acid sequence (71). Because of this, to eliminate any undetermined prediction, we decided to remove sequences from the application step where no other rule could apply as undefined (see below).

### Shannon entropy

Shannon entropy was calculated using the Entropy HIV LANL database tool (https://www.hiv.lanl.gov/content/sequence/ENTROPY/entropy_one.html)

### Prediction assessment of the model

Prediction assessment of the model was divided into training (model training and validation) and testing (application). Validation included the leave-one-animal-out cross validation, sequentially holding all the sequences from one animal out from the training set, training the classifier, and evaluating the prediction for that left out animal’s class. The leave-one-out animal approach was applied, as opposed to leave-one-sequence-out, considering that diagnosis (SIVE or SIVnoE) is assigned at the level of the subject and not at the level of individual sequences; furthermore, the genetic relatedness of the sequences derived from the same subject can lead to biased classification (19). Animals were classified as SIVE when 85% or more of their constituent sequences matched a SIVE signature. This threshold was used given that the lowest percentage obtained for a clinically diagnosed SIVE animal classified as SIVE was of 85%. The performance of the model, e.g. Accuracy, precision, were calculated only on test data.

After obtaining the model performance, we proceeded to test the original PART model (from CNS sequences), referred to as the application step, on all the remaining sequences from all animals, except the ones used for training. We divided these sequences per tissue [plasma, CD3+, CD14+, bone marrow, bronchoalveolar lavage (BAL), and lymph nodes], per time point [21, 60, 90, 180 dpi, and necropsy], and per monkey, to assess the different predictions of the model at various points in time of infection. We then proceeded to determine the exact rule used to classify each of the sequences. In this way, we intend to understand which tissue and time point could be used to predict SIVE or SIVnoE diagnosis. A total of three sequences from all sequences included in the application data sets were removed out of 3,772 sequences (0.08%). The SIVE signature that was used to classify each tissue at each time point as SIVE was also determined.

### Bayesian graphical model from amino acid sequences

The previously obtained env sequence alignment and phylogenetic tree were used for a Bayesian graphical model (BGM) analysis to see the probability of positions in the sequences co-evolving together. Fpr this, we used HyPhy with default parameters and visualized the results with the HyPhy vision BGM tool (http://vision.hyphy.org/BGM)

### HIV gp120 and SIV gp120 structural analysis

Crystal structures of gp120 were used for mapping mutations;PDB 2NY3 for HIV gp120 (38), PDB 3JCC for SIV gp120 (39). SSM in COOT (72) was used to superimpose SIV gp120 on HIV gp120, using 2NY3 as the reference structure. Interatomic distances between mutated positions and NAG (2-acetamido-2-deoxy-beta-D-glucopyranose) residues were measured in COOT. PyMOL (https://pymol.org/2/) was used to generate molecular graphic images.

### Bayesian phyloanatomy

Bayesian genealogical tree reconstruction for individual macaque-specific gp120 sequence alignments was performed using BEAST v1.8.3 (**?**) (available from http://beast.bio.ed.ac.uk/), assuming an uncorrelated relaxed molecular clock model of evolutionary rate variation across branches (**?**) and Bayesian Skyride demographic model (**?**). Prior information can be observed from the representative xml found in https://github.com/rifebd88/SIV_Phyloanatomy. A subset (500) of systematically drawn trees from the resulting posterior tree distribution for each macaque was obtained for use as empirical data for further analysis, owing to the computational complexity of integrating over the possible tree space for a large number of taxa (50). The tree subsets and script used to generate them can also be found at https://github.com/rifebd88/SIV_Phyloanatomy. Macaque-specific gp120 sequence alignments were categorized according to SIVE diagnosis (SIVE and SIVnoE) and treated as individual partitions according to a hierarchical phylogenetic model (HPM) in subsequent Bayesian analysis. An asymmetric transition rate matrix within the Bayesian stochastic search variable selection (BSSVS) allowed for inferred directionality of viral dissemination patterns occurring at significantly non-zero rates between discrete sampled anatomical compartments. Within the HPM, an epoch model (**?**) was used to infer spatiotemporal dissemination patterns during intervals of time over the duration of the evolutionary history of the viral population (assumed constant over time). Time intervals included the first 21 dpi (approximating acute infection), last 21 days prior to necropsy (approximating AIDS onset), and the duration between (approximating asymptomatic infection). Prior information can be observed from the representative xml found in https://github.com/rifebd88/SIV_Phyloanatomy. Effective Markov chain Monte Carlo sampling (**?**) for all Bayesian analyses was assessed by calculating the effective sample size (ESS) for each estimated parameter. ESS values > 200, calculated in Tracer (available from http://beast.bio.ed.ac.uk/Tracer), were considered suitable indicators of effective sampling. Bayes factor (BF) support (BF>3) (**?**) for non-zero transition rates of SIV viral lineages among discrete anatomical locations within the host was assessed at the hierarchical level using SPREAD (**?**) (latitude and longitude coordinate designations of ‘1’ for all anatomical locations).

